# Mechanistic models for West Nile Virus transmission: A systematic review of features, aims, and parameterisation

**DOI:** 10.1101/2023.10.26.564175

**Authors:** Mariken de Wit, Afonso Dimas Martins, Clara Delecroix, Hans Heesterbeek, Quirine ten Bosch

## Abstract

Mathematical models within the Ross-Macdonald framework increasingly play a role in our understanding of vector-borne disease dynamics and as tools for assessing scenarios to respond to emerging threats. These threats are typically characterised by a high degree of heterogeneity, introducing a range of possible complexities in models and challenges to maintain the link with empirical evidence.

We systematically identified and analysed a total of 67 published papers presenting compartmental West Nile Virus (WNV) models that use parameter values derived from empirical studies. Using a set of fifteen criteria, we measured the dissimilarity compared to the Ross–Macdonald framework. We also retrieved the purpose and type of models and traced the empirical sources of their parameters.

Our review highlights the increasing refinements in WNV models. Models for prediction included the highest number of refinements. We found uneven distributions of refinements and of evidence for parameter values. We identified several challenges in parameterising such increasingly complex models. For parameters common to most models, we also synthesise the empirical evidence for their values and ranges. The study highlights the potential to improve the quality of WNV models and their applicability for policy by establishing closer collaboration between mathematical modelling and empirical work.

## INTRODUCTION

West Nile virus (WNV) is a mosquito-borne pathogen that has caused outbreaks worldwide. It is transmitted between mosquitoes and birds, but the pathogen can spill over to humans and horses. Although most human cases are asymptomatic, it can cause a variety of symptoms ranging from fever to encephalitis in the most severe cases [1]. There is currently no available vaccine or specific treatment against West Nile Virus infections in humans [1]. Thus, current prevention measures mostly consist of mosquito control campaigns [2].

West Nile Virus can be transmitted to a wide range of species [3], making its dynamics complex. WNV is primarily transmitted through the bites of infected mosquitoes, with birds serving as the main reservoir host [4]. *Culex* species are the main vector that amplifies WNV as it feeds preferably on competent bird species [5]. This maintenance process in mosquito and bird populations is characterised by a high degree of heterogeneity. Within a single *Culex species*, there can be variations in biting behaviour and transmission efficiency based on factors such as age, sex, time of the year, and infection status [6, 7]. Occasionally, there is spill-over to dead-end species like humans and horses, which are incapable of further transmitting it [8].

Mathematical models aim to capture the diversity of species and processes involved in West Nile Virus transmission dynamics. These models capture dynamical processes involved in WNV transmission and thereby contribute to our knowledge of WNV and help to predict the course of future outbreaks. Mathematical models can help understand the transmission and establishment of WNV, as well as which factors contribute to this, by estimating metrics such as the basic reproduction number R_0_, the force of infection, or human infection risk [9]. Models are also used to inform policy, for example by estimating the current levels of WNV transmission based on surveillance data, by determining the risk of future outbreaks, or by estimating the effect of control interventions. This can be done on both short and long-term scales and coarse and fine spatial scales and could specifically include predictions under change scenarios. Two recent systematic reviews have studied the use of WNV models for guiding interventions and made recommendations on how to make models more useful for policy-making [9]. Both reviews recommended the development of models on finer spatial scales as that corresponds better to the scale at which vector control interventions are implemented. Keyel et al. [10] also recommended a closer alignment of model outputs with required information for decision-making.

The foundation for many modelling efforts to understand WNV dynamics was laid by Ronald Ross’ work on malaria, extended by George Macdonald [11, 12]. Although several Ross-Macdonald-type models have been developed, they typically centre around the concept of a basic reproduction number and include a simplified description of the transmission cycle [13]: 1) an infectious mosquito passes the pathogen to a host upon a bite, 2) the pathogen infects the host, multiplies and reaches high densities in the host bloodstream, 3) the pathogen is passed to a mosquito upon a bite on the infectious host, and 4) the pathogen infects the mosquito and multiplies so that the virus reaches sufficiently high concentrations in the salivary glands to be transmitted upon a bite. Some common assumptions of these models are: bites are evenly distributed among the host population, transmission only happens between vectors and hosts, the incubation period and biting rate are constant over time as is the mosquito-to-host ratio. This is implemented in the framework of compartmental models (also known as SIR-type models), in which the host and vector populations are divided into classes based on their infection status. Within this context, models have been expanded to answer present-day challenges posed by mosquito-borne diseases. Examples of adaptations include temperature dependence, vertical transmission, and multiple host species.

To study how the Ross-Macdonald approach has changed since its development, Reiner and Perkins et al. [14] compiled an exhaustive list of mathematical models for mosquito-transmitted diseases, spanning over 388 models published between 1970 and 2010. Of these, 31 had WNV as the pathogen of study. The authors argue that many models developed over the years still bear a strong resemblance to the foundational Ross-Macdonald ideas and suggest that new theory could benefit from including concepts like heterogeneous mosquito biting, poorly mixed mosquito-host encounters, spatial heterogeneity, and temporal variation in the transmission process.

More than a decade after the initial review, both the geographic range of WNV has expanded, as has the number and complexity of models aimed specifically at the WNV system. These developments raise important questions about the availability and suitability of empirical studies necessary to inform these more complex models and their application in new locations. Adding heterogeneity in models is essential, as is considering new environmental and ecological contexts for emergence and spread. Models then become more parameter rich, while at the same time, the values for parameters common to the most basic mechanistic descriptions of the system can differ between regions and contexts where they were previously determined and the new situations where the model will be applied. With an increasing use of models focused on understanding and predicting outbreaks to direct policy, these developments can have important consequences for decision-making in new areas or populations.

Building on the study by Reiner and Perkins et al., we first describe trends in assumptions of compartmental models, with a specific focus on WNV, extending the analysis with models published between 2010 and 2022. Additionally, we explore the purpose and type of WNV models, also in relation to their similarity to the basic Ross-Macdonald framework, to identify the conceptual developments that have been implemented in the last decade. We then provide insight into the evidence base of empirical studies that are cited to have informed the values of the parameters common to most of these models. Finally, we synthesise the challenges emerging from this.

## METHODS

### Literature search strategy

We searched the peer-reviewed literature for mechanistic models of WNV transmission on a population level (figure 1) that used parameter values based on data. Specifically, we were interested in compartmental models that modelled population dynamics in time with differential equations. The database search query combined terms related to “West Nile virus”, “mathematical”, and “model” (see Supplementary material 1 for full details) and used these to search PubMed, Scopus, and Web of Science. To minimise the risk of missing relevant papers, we then used a forward snowballing approach [15], where we used the 15 most cited studies identified through the database search to add the papers that cited these. The same inclusion and exclusion criteria were applied in the systematic database search and in the snowball round. The final search was completed on 6 September 2022. Inclusion and exclusion of papers were performed by two reviewers independently. Disagreements were discussed until a consensus was reached, and a third author was involved in making the final decision when necessary. The full list of identified papers, selection steps, and final list of included papers is presented in Supplementary Material 2.

**Figure 1.**
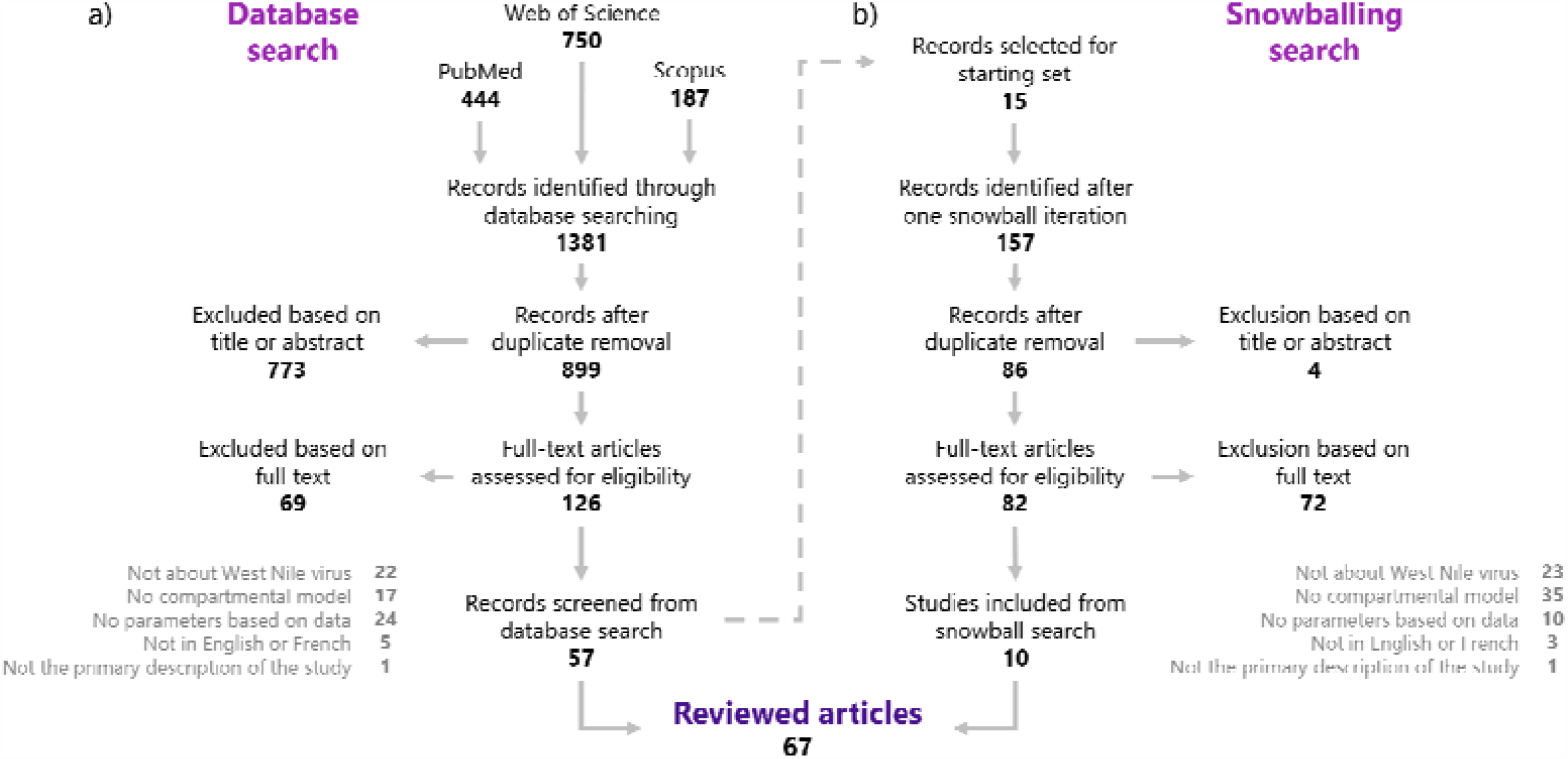
Complementing the systematic literature search with one iteration of snowballing search. **a)** PRISMA diagram depicting inclusion and exclusion steps of the database searching. **b)** Decision tree of the snowball searching.

### Model classification

For each publication, we retrieved the purpose and type of the model and classified the refinements compared to the basic Ross-Macdonald assumptions. We defined three categories for the purpose of models: understand, predict, and control, as defined in Cecilia et al. [16] Models were classified as “understand” if they aimed at exploring the impact of various mechanisms on the transmission dynamics, as “predict” if they were forecasting the evolution of WNV transmission in time and as “control” if they were exploring the impact of intervention strategies. Additionally, we used the categories defined in Cecilia et al. to classify the types of models: applied, theoretical, and grey. Applied models describe a specific area and use data to calibrate or validate the model, theoretical models are generic models and do not use any data, and grey refers to models that do not fit into these categories. We classified the refinements by calculating a dissimilarity index as proposed in Reiner and Perkins et al. [14] to quantify the divergence of the models from the basic Ross-Macdonald assumptions. For each publication, the model was classified based on 15 criteria (Table 1). For each criterion, the model scored one if it is refined compared to the assumption of the basic Ross-Macdonald model and zero otherwise. The resulting sum of the scores for all criteria is a dissimilarity index, between 0 and 15, for each publication. When several models were described in a publication, we used the one having the most refinements. Each publication was read and classified by two authors. Disagreements were discussed until a consensus was reached, and a third author was involved in making the final decision when necessary.

**Table 1.** Details on the dissimilarity index criteria.

| Index | Criterion | Ross-Macdonald assumption | Deviation from Ross-Macdonald assumption |
| --- | --- | --- | --- |
| 1 | Aquatic stages of mosquito populations | Not modelled | Explicitly modelled, with at least one state variable |
| 2 | Number of spatial locations | One without immigration | One with immigration, or more than one |
| 3 | Number of mosquito taxa, genotypes or phenotypes | One | More than one |
| 4 | Number of pathogen taxa, genotypes or phenotypes | One | More than one |
| 5 | Number of vertebrate taxa, genotypes or phenotypes | One | More than one (including non-competent hosts) |
| 6 | Mosquito mortality in the absence of control | Constant per capita mortality rate | Any refinement (temperature-dependent, humidity-dependent...) |
| 7 | Mosquito blood-feeding rate in the absence of control | Constant per capita blood-feeding rate | Any refinement (temperature dependent, dependent on host availability, ...) |
| 8 | Proportion of blood meals on hosts | Feeding on other hosts was not included | Any refinement (feeding on non-competent hosts...) |
| 9 | Pathogen latency in mosquitoes | Not modelled | Explicitly modelled as a state variable |
| 10 | Waning immunity | Not modelled | Explicitly modelled |
| 11 | Superinfections and co-infections | Not modelled | Explicitly modelled |
| 12 | Distribution of blood meals among hosts | Homogeneous distribution of bloodmeals among vertebrate hosts | Heterogeneous distribution of bloodmeals among vertebrate hosts |
| 13 | Mixing | Well-mixed | Not well-mixed, modelled using a contact network or individual-based model |
| 14 | Transmission from mosquito to host | Constant | Any refinement (temperature-dependent, time-dependent...) |
| 15 | Transmission from host to mosquito | Constant | Any refinement (temperature-dependent, time-dependent...) |

### Empirical evidence for parameter values

To study the use of available evidence from empirical studies in model parameterisation, we extracted cited references for parameter values and traced the empirical sources of these values. We focused on values for the six most common virus-specific parameters: extrinsic incubation period (the time the virus takes to incubate within the vector), intrinsic incubation period (the time the virus takes to incubate within the host), recovery rate (i.e., duration of infectious period), disease-induced mortality rate, transmission probability from vector to host and from host to vector. We extracted references provided for these parameter values from all studies and traced back to empirical studies underlying these values (e.g., in case the cited reference was not an empirical study but a modelling paper or a review). For each of the empirical studies, we counted the number of times it was used as underlying source for parameter values by studies included in this review.

## RESULTS

We identified a total of 67 papers that published compartmental WNV models using parameter values based on data [17–83]. The oldest paper included was published in 2001 and since then, the number of modelling papers published per year has increased over time (R=0.59, p-value=7.5 * 10^-3^, figure 2). The majority of studies included in our review were published in the past decade (65%), with 36% being published in the past five years.

**Figure 2.**
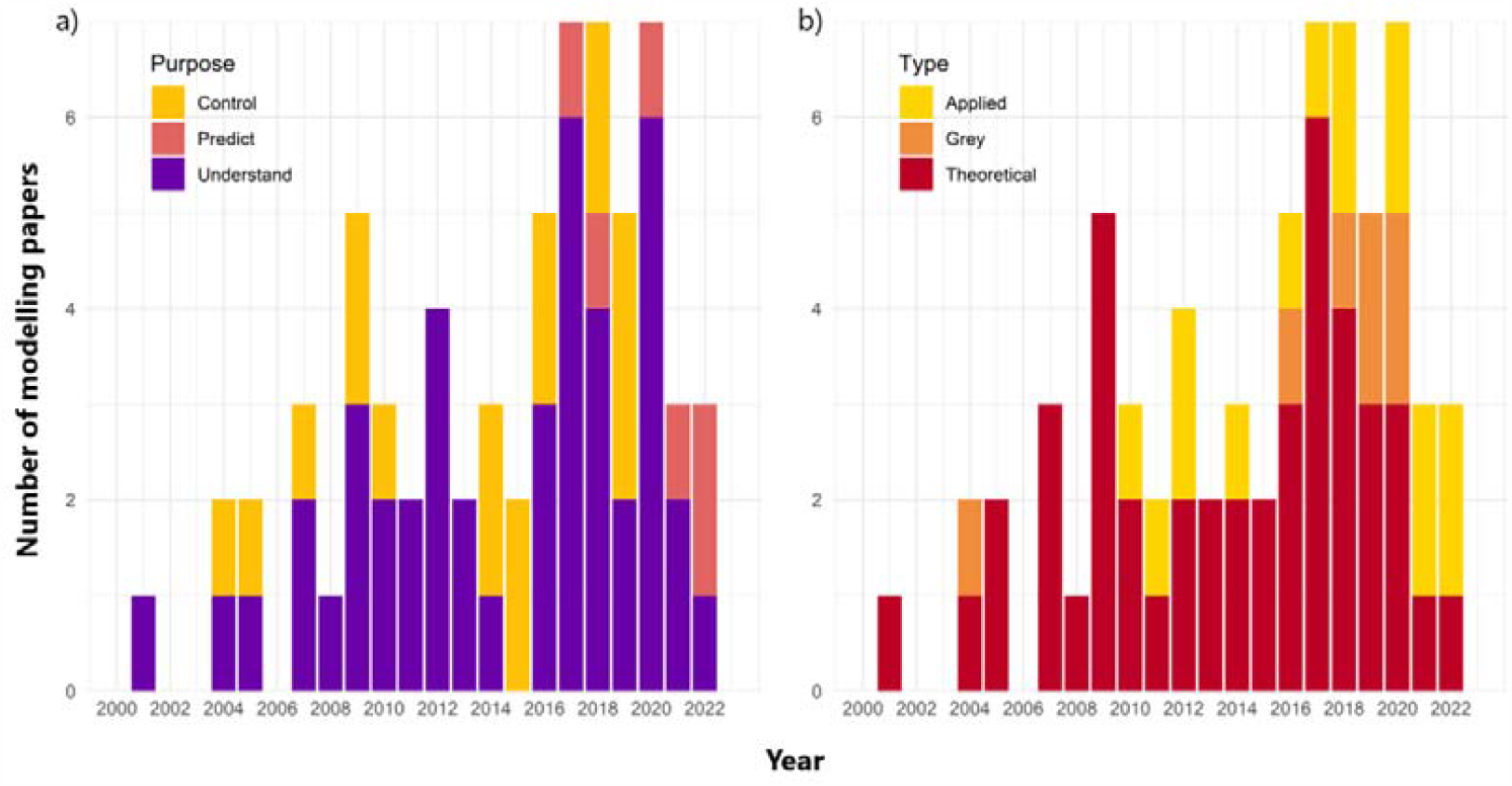
Frequency of modelling studies included in the review classified according to **(a)** their research purpose and **(b)** study type.

The dissimilarity index, calculated using 15 criteria (Table 1), increased slightly over time (R=0.27, p-value=0.026), indicating that WNV compartmental models tended to increasingly deviate from the approaches of Ross and Macdonald (figure 3a). The mean number of refinements included was 3.1. We identified 66% of models (n=44) aiming at understanding transmission, 25% (n=17) aiming at evaluating control strategies and 9% (n=6) aiming at predicting case numbers. Models aiming at predicting had a higher dissimilarity index on average (mean=6.3, p-value=2.1*10^-5^). Additionally, 67% of the models (n=45) were classified as theoretical, 22% (n=15) as applied and 10% (n=7) as grey. Applied models had a higher dissimilarity index on average (mean=5.1, p-value=1.1*10^-5^). A variety of data types were used in applied and grey models, including demographic and environmental data such as mosquito abundance and temperature, as well as epidemiological data such as incidence in humans, serological data, and prevalence in trapped mosquitoes.

**Figure 3.**
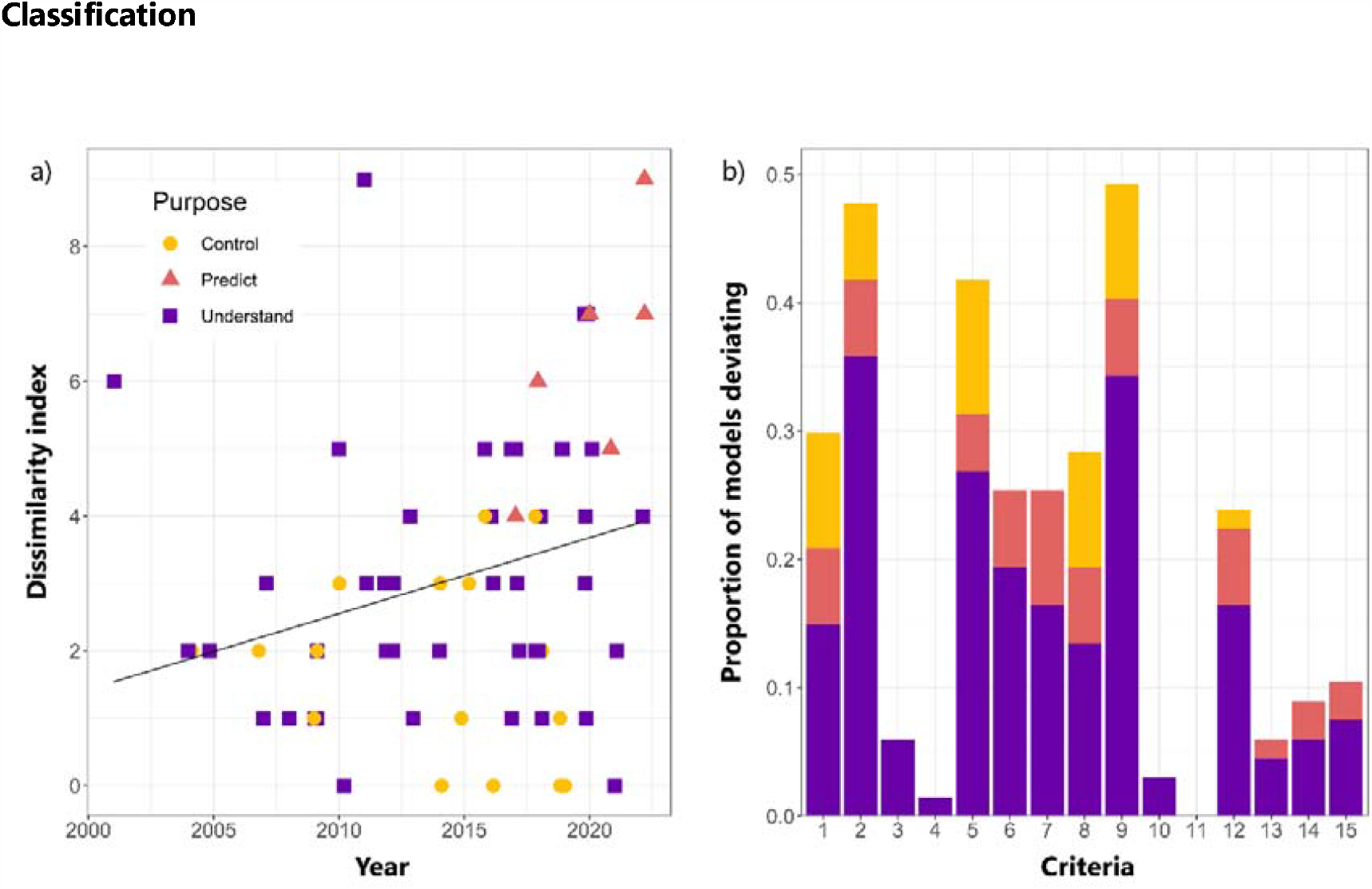
(a) Dissimilarity index of publications per year of publication. **(b)** Proportion of models deviating from criteria of the dissimilarity index.

The most common refinements of the Ross-Macdonald framework were including multiple spatial locations (or taking migration into account) (48%), including pathogen latency in mosquitoes (49%), and including multiple host species (42%) (figure 3b). The majority of models with multiple host species (n=15, 53% of those with more than one host species) included non-competent hosts (humans n=14, equids n=1). Additional competent hosts were studied in 16 studies (57% of those with more than one host species studies), including up to 8 different bird species. Contrarily, only 6% of models included more than one mosquito species, of which 75% included multiple *Culex* species, and one included *Aedes albopictus* in addition to the most commonly modelled *Culex pipiens*. The least common refinements were waning of immunity (3%), number of pathogen taxa (1%), and superinfection (0%). The only study considering more than one pathogen included avian malaria in addition to WNV but did not explicitly model co-or superinfection [33]. All classification results per paper are presented in Supplementary Material 3.

### Empirical underpinnings of key model parameters

The reviewed body of literature resulted in a collection of infection and transmission parameters. The complete datasets on the parameters and their details can be found in the Supplementary Materials 4. Here, we describe the body of evidence used by the authors of the studies to estimate key virus-specific parameters: the disease-induced death rate, the recovery rate, the intrinsic incubation period, the latency or extrinsic incubation period, the transmission probability from host to vector, and the transmission probability from vector to host.

The number of unique empirical studies used as underlying source for parameters varied from nine for the intrinsic incubation rate to 24 for the transmission probability from mosquito to host. Some papers cited these empirical studies directly, but most (63%) used other models or reviews as a reference (i.e., indirect citation, figure 4). A small number of papers were clearly used more frequently, while 49% of studies were only used once. This trend was especially strong for the recovery rate parameter, with one paper [84] used as a source in 73% of all citations to empirical studies. A comparison of the parameter values used in the models for the most cited empirical sources is provided in Supplementary Materials 6.

**Figure 4.**
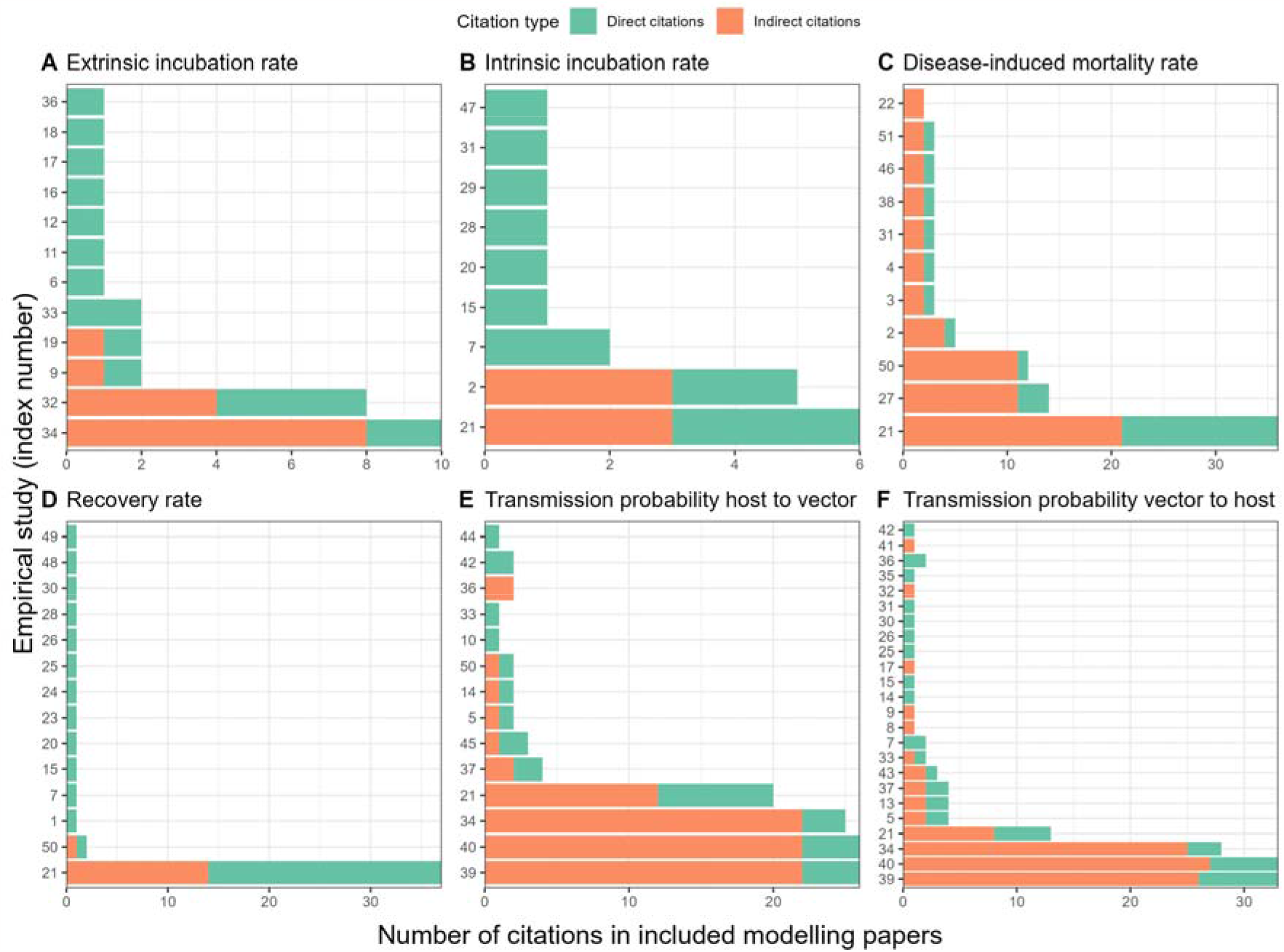
(a-f) Number of direct (green) and indirect (orange) citations per experimental study used as source for each key model parameter. Full list of original sources (index numbers on y-axis) and model studies that cited them can be found in the Supplementary Material 5.

## DISCUSSION

We identified 67 papers presenting compartmental WNV models. The number of refinements to the Ross-Macdonald framework increased over time. Models aiming at predicting transmission and those including data from a specific area included the highest number of refinements. Compared to Reiner and Perkins et al. [14], models included here addressed more refinements on average. We found an uneven distribution in which model refinements are addressed, with spatial structure and/or migration and pathogen latency in mosquitoes being addressed in about half the models, while others, such as superinfection, were never addressed. Most papers referred to other modelling papers and/or reviews rather than the underlying empirical studies for their parameter values. A small number of empirical studies were highly influential through indirect citations and frequently used as a source of parameter values, whereas half of the empirical studies used were only referred to once.

One of the main recommendations put forward in Reiner and Perkins et al. [14] related to the incorporation of host heterogeneity, which we encountered in several WNV models. This was achieved through the inclusion of multiple host species with host-specific infection parameters in 28 models (42%) ([20, 70] and/or species-specific biting preferences in 16 models (24%) (e.g.,[47, 58]) sometimes extended with changes in biting preferences over time in 3 models (4%) [28, 29, 44]. Like Reiner and Perkins et al., we observed that heterogeneity through imperfect mixing of hosts and mosquitoes and any spatial heterogeneity was still rarely included. A major limitation in the inclusion of refinements is the availability of empirical data to underpin parameterisation. This is illustrated by the development of models including multiple host species. The experimental work by Komar et al. [84] enabled researchers to study the role of multiple host species and the differences between these on WNV dynamics. All papers including multiple host species were published after this experimental study. Especially models classified as ‘predictive’ and ‘applied’ would benefit from high-quality empirical studies as these types of models included most refinements and relied on data for their modelling efforts.

### Challenges in translating parameter values from empirical sources

Based on our results, we observed two trends regarding the use of empirical evidence in modelling efforts. We found that the body of evidence from empirical studies is not used to its full extent with many studies rarely being used and a small number of studies being used frequently mostly because they were cited in other model papers. Secondly, we noticed that models using parameters based on data (the focus of this review) are limited in what refinements can be studied by the availability of reliable and relevant data. The inclusion of model refinements often leads to an increase in the number of model parameters, giving rise to challenges in model parameterisation. Based on how empirical studies were used in the model papers, we highlight three challenges that researchers encounter when parameterising their models with empirical data: A) experimental conditions may not reflect conditions in the modelled system, B) not all model parameters can be (easily) estimated in an experimental setting, C) it is not straightforward to combine results from multiple studies, especially when different species and pathogen strains are involved.

One of the major assumptions that must be made is that **experimental conditions reflect conditions in the modelled system**. This includes environmental conditions such as temperature and humidity as well as inoculation route and dose. Several steps of the transmission cycle have been shown to be sensitive to temperature conditions [85]. Only ten of sixty-seven included models used temperature-dependent parameters. However, instead of parameter values changing as a function of temperature, models can also use temperature-explicit parameters with constant values, which represent a specific temperature scenario, such as in Vogels et al. [61]. The extrinsic incubation period strongly depends on temperature [86] so when interpreting mosquito infection experiments, it is important to compare the temperature in the experiment to the temperature of the modelled system to determine if the results from this experiment are useful. This is, however, complicated by the fact that temperature can be controlled and kept constant in laboratory settings, whereas it is difficult to know what temperatures mosquitoes are exposed to outside an experimental setting. Although coarse-scale air temperature is often used as a proxy, these estimates can differ significantly from the small-scale microclimates mosquitoes are exposed to, which can have a substantial impact on transmission potential [87]. Only one of the cited empirical studies for EIP looked at the impact of varying temperatures on EIP (by using an outside cage) [88]. Also, the inoculation route and dose have a strong impact on experimental results. In the two most cited papers as a source for the host-to-mosquito transmission probability, mosquitoes fed on viraemic chickens with two different viremia levels, showing that the exposure dose has a large impact on resulting transmission rates [89, 90]. After selecting a relevant route that matches the modelled system, choosing which dose to use for the parameter value can be difficult as it is not always known which best reflects natural infection, especially as viremia levels change over the course of an infection. Ideally, transmission rates are determined for animals that were exposed through a natural route. For WNV, this was done through a host-mosquito-host system [91], and this has also been done using a host-mosquito-host system as well as mosquito-host-mosquito system for Rift Valley fever virus [92, 93].

### It is not always possible to directly estimate model parameters in experimental and field studies

Additionally, the quantities estimated in experimental and field studies do not always match the interpretation of model parameters. Experimental infection studies investigating the extrinsic incubation rate, included in 49% of the included studies, often show viral titres over time after infection, but interpreting which titre corresponds to a dose sufficiently high to represent infectiousness is not straightforward. Infectiousness of both host and mosquitoes is a function of viral load, with a higher viral load corresponding to a higher transmission probability [94]. Additionally, individuals with low viral titres can also contribute significantly to the transmission dynamics, depending on their numerical prominence [95, 96]. Defining a cut-off value for the extrinsic incubation period is therefore a somewhat arbitrary choice. Secondly, heterogeneous biting of the mosquito towards different hosts was accounted for in 24% of the models using feeding preference coefficients. These are typically quantified in field experiments, where captured blood-fed mosquitoes are analysed to determine what hosts they fed on [58, 97]. However, these estimates are limited to the context of host availability in the field settings [98]. Relative host availability is known to influence mosquito feeding behaviour [99] but mosquito host-seeking behaviour operates on small spatial scales for which host availability is often unknown. Thus, results of feeding preference experiments cannot be easily translated into a feeding preference coefficient in a model and context with different host availability. This challenge may limit the integration of heterogeneous biting in models. This limitation hampers, among others, the understanding of the impact of the ecosystem composition or the change of biodiversity on disease risk using models.

### It is sometimes required to combine multiple empirical studies to parameterise a model

When multiple estimates of a parameter are available, combining studies can increase the accuracy of parameter values or provide a better notion of the variability of the parameter. Similarly, when no information is available for a species of interest, multiple studies can be combined to average parameter values of related species [44]. However, studies can differ in their designs and outcome measures, and each have their own limitations, such as measurement errors, biases, or missing values, which can introduce uncertainty into the parameter estimation process. Additionally, different data sources may provide conflicting or inconsistent information, requiring careful consideration and potential reconciliation of the discrepancies. For example, the empirical studies used for parameterising the disease-induced death rate in included models show considerable variation in mortality rates in crows. Laboratory experiments reported a 100% mortality rate [100], while field observations indicated an overall mortality of 43% [101], with further variations being observed in different locations [41, 102, 103]. While for some species multiple studies are available, authors including less well-studied species may not be able to find empirical evidence for their species of interest. For example, in Lord et al., the authors had no available information for their species of interest and so decided to consider a large interval for those parameters based on sources for similar species [44].

Modelers are often confronted with multiple of the described challenges when developing realistically parameterised models. In such cases, several approaches exist to ensure that model outcomes reflect the uncertainty in the choice of model parameters. In the reviewed literature, some authors accounted for parameter uncertainty by using ranges rather than single values for parameters, such as in [44]. When running the model several times, taking samples from this range, the outcome measures can be presented as an interval rather than a single value. This helps capture the inherent uncertainty and provides a range of possible model outcomes. Another approach to this is to use probabilistic methods, like Monte Carlo simulations, to sample parameter values from probability distributions. Alternatively, explicit sensitivity or elasticity analyses can be performed to assess how variations in parameter values affect model outcomes like the basic reproduction number R_0_. Examples of this in the reviewed literature include [20, 36]. By systematically varying these parameters within plausible ranges, researchers can gain insights into the model’s robustness and identify which life-history effects have the most significant influence on WNV transmission. Ultimately, combining these approaches helps produce more reliable predictions, reflective of the uncertainty in parameter values.

There is a growing interest in comparing models and combining insights across different modelling approaches. Several projects have been undertaken in which public health questions were addressed by combining model-based outcomes in an ensemble forecast [104–106]. While guidelines for combining empirical studies exist (e.g., Consolidating Standards of Reporting Trials (CONSORT) and Strengthening the Reporting of Observational Studies in Epidemiology (STROBE)) and are widely used, such methods for systematically reviewing, comparing and combining model studies are still being developed [107–109]. As Pollet et al. [107] highlighted, such guidelines have the potential to improve the quality and useability of model-based prediction for public health. Approaches to evaluate and synthesise results across model studies need to address high-level questions such as how to bring together insights from theoretical models with detailed data-driven simulation models and what determines the quality of a model, as well as more practical questions such as how to synthesise results across different spatial and temporal scales. Additionally, this could help understand how including specific refinements in a model affects the outcome. Such an endeavour for WNV holds potential, not only in improving modelling predictions but also in assisting the establishment of policy guidelines for more efficient control of the disease.

## CONCLUSION

In conclusion, transmission models of WNV have recently increasingly deviated from the basic Ross-Macdonald framework, which allowed for more complex questions to be answered. However, some extensions have received more attention than others, as they are dependent on available parameter estimates. Especially applied, predictive models included a large number of refinements. This implies that these types of models, the ones often used to answer policy-related questions, are particularly sensitive to the availability and use of empirical studies to inform parameterisation. Bridging the gap between empirical data and mathematical modelling presents its own set of challenges. Translating parameter values from empirical sources into models can be a complex task, as it requires careful consideration of various factors. It is important that experimental conditions reflect the conditions in the modelled system as best as possible to ensure meaningful and reliable results. However, estimating model parameters directly from experimental and field studies is not always feasible due to practical limitations and constraints. Furthermore, the integration of multiple empirical studies to parameterise a model poses additional challenges, such as inconsistencies and variations in data sources and methodologies. Overcoming these obstacles requires a collaborative effort between mathematical modellers and experimentalists, where both are involved in the design of empirical studies and in the design and parameterisation of models. This will open up opportunities to advance our understanding of WNV transmission and enhance its practical applications in decision-making for policy.

## SUPPLEMENTARY MATERIALS

### Supplementary Material 1: Systematic database search routine

The file describes the search strategy used to identify relevant papers.

### Supplementary Material 2: Paper selection

This file provides an overview of all papers identified in the search, the results of the selection steps and a final list of included papers.

### Supplementary Material 3: Classification

This file provides the results of the classification for each included paper. This contains both the purpose of models and the refinements.

### Supplementary Material 4: Parameters

This file presents the full list of parameters collected from papers included in this review. Parameters are organised in two categories: transmission parameters and infection parameters. The tabs contain the following information about each parameter: notation used; description/interpretation provided by the article; value/estimate/mean, lower and upper estimates; units; vector and host species considered; if it is temperature dependent or not; and a formula when given; whether the value was based on literature, assumed, or estimated by comparing the model output to data. The index number of paper refers to the list of included papers in this review as presented in Supplementary Material 2.

### Supplementary Material 5: Original sources

This file presents the list of empirical studies used as underlying sources for parameter estimates used in the included model papers. Index numbers correspond to the y-axis in figure 4. For each of the six main parameters we show which empirical studies were used by the models and whether these empirical studies were cited directly (i.e., direct reference to this study) or indirectly (i.e., paper cites another paper which ultimately references to this study).

### Supplementary Material 6: Variation in parameters

This file presents an analysis of the parameter values used in models which all used the same, most frequently cited, empirical study as a source.

