## Supplemental 1 for "Mechanistic models for West Nile Virus transmission: A systematic review of features, aims, and parameterisation"

***Supplementary Material 1: Systematic database search routine***

The following queries were used in our systematic database search:

- Pubmed
  - *((“models, theoretical”[MeSH Terms] OR “mathematic*” [Title/Abstract] OR “simulati*”[Title/Abstract] OR “mechanistic”[Title/Abstract] OR “compartment*”[Title/Abstract] OR “sir”[Title/Abstract] OR “differential”[Title/Abstract] OR “deterministic”[Title/Abstract] OR “theoretical”[Title/Abstract]) AND (“west nile virus”[MeSH Terms] OR “west nile*”[Title/Abstract] OR “egypt 101*”[Title/Abstract] OR “kunjin*”[Title/Abstract])) AND (model*[Title/Abstract])*
- Web of Science
  - *((TS=(West Nile* OR Egypt 101* OR Kunjin*)) AND TS=(mathematic* OR simulat* OR mechanistic OR compartment* OR SIR* OR differential OR deterministic OR theoretical)) AND TS=(model*)*
- Scopus
  - *TITLE-ABS-KEY("west nile*" OR "egypt 101*" OR "kunjin*") AND TITLE-ABS-KEY("mathematic*" OR "simulat*" OR "mechanistic" OR "compartment" OR "SIR" OR "differential" OR "deterministic" OR "theoretical") AND TITLE-ABS-KEY("model*")*
