## Supplemental 6 for "Mechanistic models for West Nile Virus transmission: A systematic review of features, aims, and parameterisation"

***Supplementary Material: Variation in parameters estimates based on the most cited empirical study***

Figure S1 shows parameter estimates used in the models citing the most cited empirical study. These empirical studies are:

- Extrinsic incubation rate: Sardelis et al. 2001 [1]
- Intrinsic incubation rate: Komar et al. 2003 [2]
- Disease-induced death rate: Komar et al. 2003 [2]
- Recovery rate: : Komar et al. 2003 [2]
- Transmission host to vector: Turel et al 2001 [3]
- Transmission vector to host: Turel et al 2001 [3]

Some of these examples apply estimates to a wide range of different species, while citing the same source. For instance, Maidana and Yang [4] estimated the recovery rate for 26 different species, resulting in the large interval of (0.18-3.3) seen in the plot.

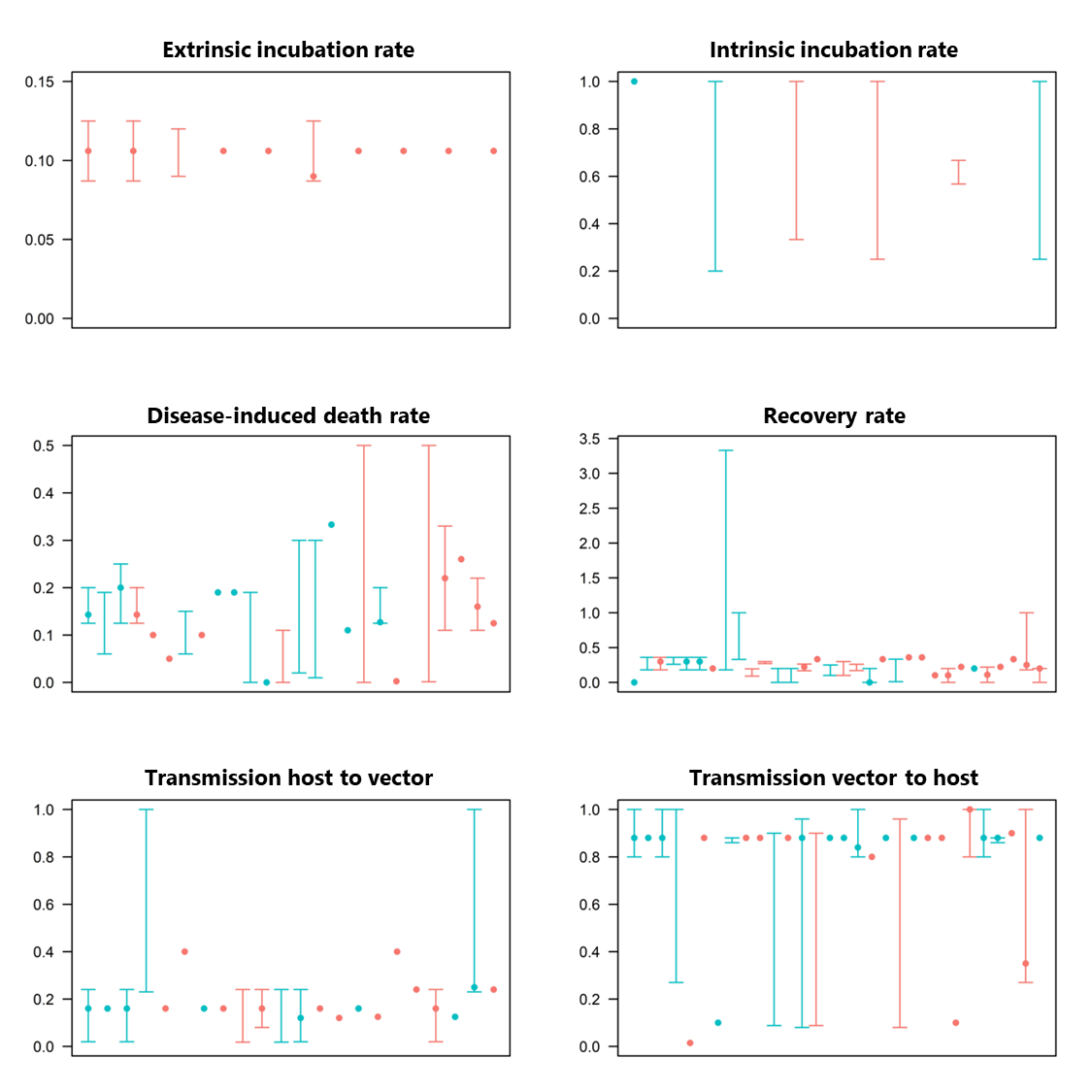

**Figure S1** Parameter values used in model papers (from oldest to most recent) citing the same experimental study. Models using estimates from the original sources in different or unspecified species are plotted in red and models applying the estimates to the same species are plotted in blue. Results are shown for model papers citing the most frequently cited experimental study for each parameter. Dots represent point estimate and bars the lower and upper bounds of intervals considered.

1.	Sardelis MR, Turell MJ (2001) Ochlerotatus j. japonicus in Frederick County, Maryland: discovery, distribution, and vector competence for West Nile virus. J Am Mosq Control Assoc 17:137–41

2.	Komar N, Langevin S, Hinten S, Nemeth N, Edwards E, Hettler D, Davis B, Bowen R, Bunning M (2003) Experimental Infection of North American Birds with the New York 1999 Strain of West Nile Virus - Volume 9, Number 3—March 2003 - Emerging Infectious Diseases journal - CDC. Emerg Infect Dis 9:311–322

3.	Turell MJ, O’Guinn ML, Dohm DJ, Jones JW (2001) Vector competence of North American mosquitoes (Diptera: Culicidae) for West Nile virus. J Med Entomol 38:130–134

4.	Maidana NA, Yang HM (2009) Spatial spreading of West Nile Virus described by traveling waves. J Theor Biol 258:403–417
